# The influence of temporal predictability on express visuomotor responses

**DOI:** 10.1101/2020.08.28.269449

**Authors:** Samuele Contemori, Gerald E. Loeb, Brian D. Corneil, Guy Wallis, Timothy J. Carroll

**Affiliations:** Centre for Sensorimotor Performance, School of Human Movement and Nutrition Sciences, The University of Queensland, Brisbane, Australia; Department of Biomedical Engineering, University of Southern California, Los Angeles, California, USA; Department of Physiology and Pharmacology, Western University, London, Ontario, Canada; Department of Psychology, Western University, London, Ontario, Canada; Robarts Research Institute, London, Ontario, Canada

**Keywords:** rapid visuomotor responses, visually-guided reaching, stimulus timing prediction, electromyography, moving target, transient visual stimulus

## Abstract

Volitional visuomotor responses in humans are generally thought to manifest 100ms or more after stimulus onset. Under appropriate conditions, however, much faster target-directed responses can be produced at upper limb and neck muscles. These “express” responses have been termed stimulus-locked responses (SLRs) and are proposed to be modulated by visuomotor transformations performed subcortically via the superior colliculus. Unfortunately, for those interested in studying SLRs, these responses have proven difficult to detect consistently across individuals. The recent report of an effective paradigm for generating SLRs in 100% of participants appears to change this. The task required the interception of a moving target that emerged from behind a barrier at a time consistent with the target velocity. Here we aimed to reproduce the efficacy of this paradigm for eliciting SLRs and to test the hypothesis that its effectiveness derives from the predictability of target onset time as opposed to target motion *per se*. In one experiment, we recorded surface EMG from shoulder muscles as participants made reaches to intercept temporally predictable or unpredictable targets. Consistent with our hypothesis, predictably timed targets produced more frequent and stronger SLRs than unpredictably timed targets. In a second experiment, we compared different temporally predictable stimuli and observed that transiently presented targets produced larger and earlier SLRs than sustained moving targets. Our results suggest that target motion is not critical for facilitating the expression of an SLR and that timing predictability does not rely on extrapolation of a physically plausible motion trajectory. These findings provide support for a mechanism whereby an internal timer, probably located in cerebral cortex, primes the processing of both visual input and motor output within the superior colliculus to produce SLRs.

## INTRODUCTION

Sudden events demand rapid motor responses, for example to catch an object accidentally knocked from a table, to react to an opponent in sporting context, or for self-defence to an unexpected physical threat. Often, the surrounding contextual features provide information to anticipate the timing of the event but not its exact location. For instance, when observing a child moving in an unstable way, it is possible to predict if and when a threatening event will happen in order to be prepared to react rapidly (e.g. intercepting an object knocked by the child before it hits the ground; grasping the child’s hand before it gets in contact with dangerous surfaces or objects).

Rapid visuomotor reactions have been studied in oculomotor responses (Dorris et al. 1997), neck muscles (Goonetilleke et al. 2015), arm muscles (Pruszynski et al. 2010), and leg muscles (Weerdesteyn et al. 2004). The term “express” was originally adopted by Fischer and Boch to describe the extreme short-latency at which monkeys can produce stimulus-driven saccades (i.e. less than 110-120ms from stimulus onset time; Fischer and Boch 1983; Pare and Munoz 1996; Dorris et al. 1997). Akin to the express saccade latency, the target-directed EMG activity in neck and proximal arm muscles can encode the target location within ∼100ms from the stimulus presentation (Corneil et al. 2004, 2008; Pruszynski et al. 2010). Our intention in using the term “express visuomotor responses” in the title of the paper is to draw a parallel between express saccades and the most rapid visuomotor responses that can be generated in other body parts, potentially through a common retino-tecto-reticular pathway. We hope that the use of such terminology will contribute to more widespread consideration of the implications of subcortical visuomotor transformations for theories on human motor control and behaviour.

Two different acronyms have been coined for these short latency responses in neck and arm muscles: (i) stimulus-locked responses (SLRs); (ii) rapid visual responses (RVR). The term RVR was adopted by Glover and Baker (2019) as indicative of the type of stimulus (i.e. visual) that was used to elicit the response. The term SLR is descriptive of a characteristic attribute of these EMG responses, which are consistently more time-locked (within ∼100ms) to the stimulus onset time than to reaction time, defined as initiation of the reaching movement (Corneil et al. 2004, 2008; Pruszynski et al. 2010; Wood et al. 2015; Goonetilleke et al. 2015; Gu et al. 2018; Gu et al. 2019; Atsma et al. 2018; Kozak et al. 2019). In this paper, we will use the ‘SLR’ acronym because it describes how we define and quantify the response (see materials and methods), while noting that it may be misleading as to functionality and mechanism.

The latencies of express saccades and SLRs are consistent with the minimum time that is needed to accomplish the sensorimotor transformation of visual information (Boehnke and Munoz 2008). These responses invariably reflect the stimulus location rather than the volitional movement that is ultimately produced, when these are dissociated in the anti-saccade (reviewed by Coe and Munoz 2017) and anti-reach tasks (Gu et al. 2016). The general idea is that anti-tasks require longer cortical pathways to compute the desired direction whereas express outputs are mediated at the subcortical level via the midbrain superior colliculus, whose outputs are driven directly by the stimulus location itself. For SLRs, outputs from the superior colliculus have been proposed to be delivered to spinal interneurons and motoneurons via the tecto-reticulo-spinal pathway (Corneil and Munoz 2014).

Previous work showed that SLRs are not detected consistently in all individuals. Pruszynski et al. (2010) failed to detect SLRs from surface EMG recordings and detected positive SLRs in only 7 out of 16 participants (∼44% prevalence rate) using intramuscular electrodes. Recently, Kozak et al. (2020) obtained a 100% SLR detection rate among a sample of five individuals by adopting an *emerging target paradigm*, which was motivated by earlier oculomotor studies (for review see Fiehler et al. 2019). This suggests that the circuit responsible for SLRs can be biased according to task conditions. A better understanding of the efficacy of behavioural tasks and stimuli to generate SLRs would improve both the design of experiments and the identification of the likely pathways and mechanisms that mediate and modulate them.

In the emerging target paradigm, the stimulus initially fell toward a visual barrier and re-emerged beneath it at a time that was specified by the target velocity. This made the target presentation time predictable via extrapolation of the target trajectory behind the barrier (Kozak et al. 2020). Previous studies showed that the collicular response to visual stimuli is more vigorous with moving than static stimuli (Schneider and Kastner 2005; Lau et al. 2011), raising the possibility that target motion facilitates SLRs because it provides a higher sensory salience in the superior colliculus itself. However, temporal predictability of a stimulus itself facilitates express motor outputs and could be computed outside of the superior colliculus. Both express saccades and SLRs are facilitated by the insertion of a constant and predictable time gap between a warning stimulus (e.g. offset of the fixation) and the imperative stimulus (e.g. target presentation) to move, such as in the *gap task* paradigm (Fischer and Boch 1983; Pruszynski et al. 2010; Wood et al. 2015; Glover and Baker 2019). We hypothesized that the SLR-facilitation effect of the emerging target paradigm is attributable, at least in part, to the temporal predictability of the stimulus, rather than to target motion *per se*.

In experiment 1, we used a constant velocity moving target paradigm, akin to that used by Kozak et al. (2020), thus making the stimulus onset time predictable via an antiderivative computation of its velocity (Senot et al. 2003; Zago et al. 2009; Fiehler et al. 2019), and a consequent extrapolation for a physically plausible target trajectory. In the second experiment, we kept the target onset time constant but varied the location at which it emerged beneath the barrier. This allowed us to test the effects of target location on the expression of SLRs, as well as whether a physically plausible motion trajectory helps facilitate SLRs. Furthermore, we used either sustained moving or transient flashing static targets to determine how the SLR is modulated by the temporal attributes of the visual stimuli.

We found that the presentation of predictable and transient stimuli facilitated the expression of SLRs, regardless of where the target appeared. Our findings suggest that the emerging target paradigm promotes SLRs by allowing the precise initiation of an internal timer for a learned duration, rather than by extrapolating a plausible target motion trajectory. This would be consistent with a priming effect of temporal expectations transmitted to the SLR circuit, potentially through known cortical projections to the superior colliculus (Boehnke and Munoz 2008). Furthermore, our results suggest that the SLR-circuit is sensitive to the temporal attributes of the visual stimulus and not to the vertical location of the target within a hemi-field, at least within the range of vertical visual angles explored in this experiment.

## MATERIALS AND METHODS

### Participants

Seventeen adults were recruited for this study. Fifteen subjects (11 males, 4 females; mean age: 29.0 years, SD: 5.8) participated in the first experiment. Nine of these people also participated in the second experiment, as part of a full sample of 11 adults (8 males, 3 females; mean age: 29.3 years, SD: 9.7). All participants were right-handed, had normal or corrected-to-normal vision, and reported no current neurological, or musculoskeletal disorders. All participants provided informed consent and were free to withdraw from the experiment at any time. All procedures were approved by the University of Queensland Medical Research Ethics Committee (Brisbane, Australia) and conformed to the Declaration of Helsinki.

### Apparatus

Participants were seated with the right elbow and forearm resting on a custom-built air sled that moved with limited friction on a table. The height of the chair was adjusted to allow reaching movements by the shoulder in the transverse plane centred at ∼90° of flexion (Figure 1). Both wrist and elbow mobility were restricted by orthopaedic braces. The head was stabilized by a chin and forehead rest. A constant lateral load of ∼5N was applied in the direction of shoulder transverse extension via a weight and pulley, thus increasing the baseline activity of shoulder transverse flexor muscles, including the clavicular head of the pectoralis major muscle (Gu et al. 2016). All stimuli were created in Matlab using the Psychophysics toolbox (Brainard 1997; Pelli 1997), and were displayed on a LCD monitor with a 120Hz frame rate positioned in front of the subject. The eye-to-monitor distance was ∼57cm, so 1cm on the screen corresponded to ∼1 degree of visual angle (dva). The target was a filled black circle (dimension: ∼2dva in diameter; luminance: ∼0.3 cd/m^2^) presented against a grey background (∼170.4 cd/m^2^). The aim was to create a salient and high-contrast target that has been shown to evoke short-latency collicular responses (Marino et al., 2012) and facilitate the expression of SLRs (Wood et al. 2015). The luminance of the background and target was measured with a Cambridge Research System ColorCAL MKII colorimeter. A photodiode was attached to the left bottom corner of the monitor to detect a secondary light that was presented coincidentally with the time of appearance of the real target. This allowed us to index the time point at which the stimulus was physically displayed on the screen, thus avoiding uncertainties in software execution and raster scanning of the monitor.

**Figure 1:**
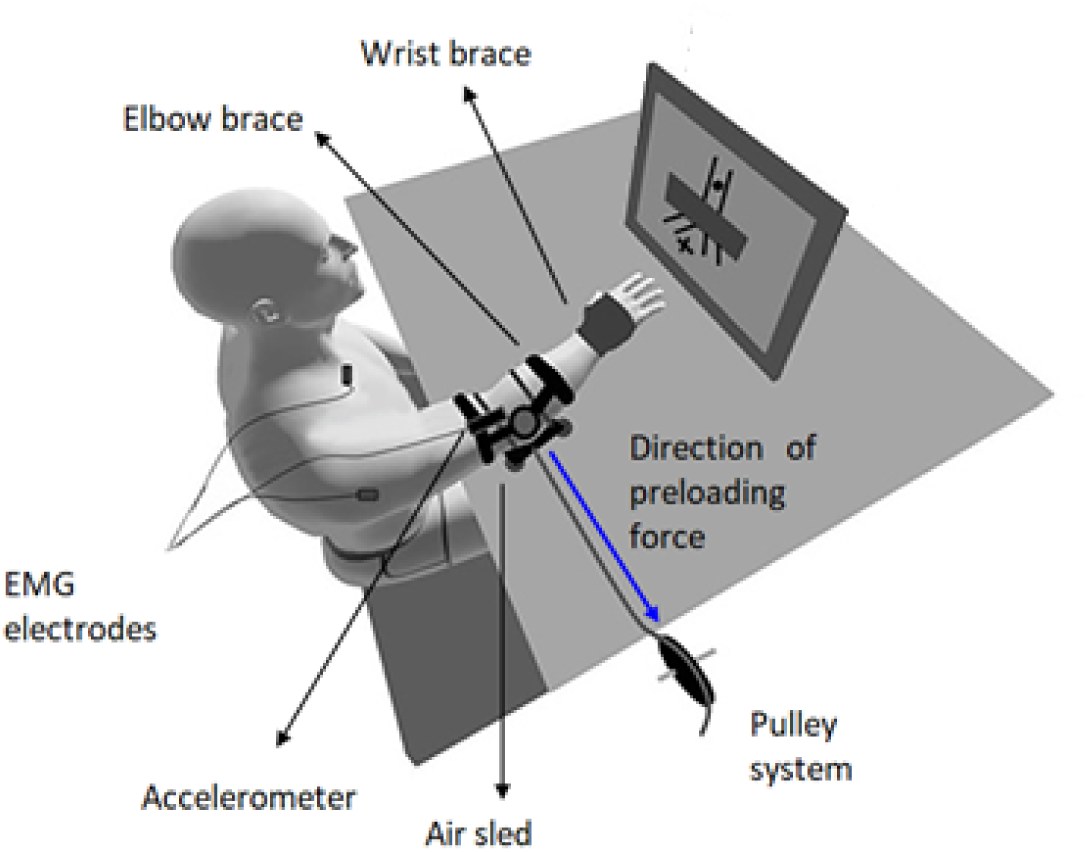
Experimental set-up. Participants sat in the experimental apparatus with their hand aligned with the fixation spot (cross bars beneath the barrier) and moved it toward the target that appeared beneath the barrier, either at the right or left of the fixation spot. The fingertip was close to the screen so that hand position corresponded closely to fixation point. Participants’ fingertips never made contact with the screen. The eye-to-monitor distance was ∼57cm, so 1cm on the screen corresponded to ∼1 degree of visual angle. Head position was stabilized by chin and forehead rests (not shown here).

### Experimental design

#### Experiment 1: predictable vs unpredictable stimulus onset time

The participants performed visually guided reaches, always starting from a constant and static upper limb position and moving as rapidly as possible toward visual targets that appeared randomly either to the right (extensor-ward) or to the left (flexor-ward) of the fixation spot, where the gaze and hand started for each trial. There were four task conditions in which the target onset time could be either predictable or unpredictable. In predictable tasks, the target was constrained to fall within an inverted y-shaped track, and a visual barrier occluded the junction point where targets could deviate left or right (Figure 2, top panel). Predictable targets dropped at constant velocity of ∼35dva/s until they passed behind the barrier, before (I) re-appearing either just beneath the barrier (7dva of fixation-target eccentricity) and continuing toward the interception point, or (II) suddenly flashing (one single flash of ∼8ms of duration) at the interception point (10dva of fixation-target eccentricity) at a time consistent with the target speed. In condition I the target appeared just underneath the barrier at ∼540ms from the trial start (onset time of target drop) and was occluded for ∼380ms. In condition II, the target appeared transiently at the interception point at ∼720ms from the trial initiation and remained invisible for ∼560ms. Unpredictable targets disappeared from the origin and suddenly reappeared at a random time underneath the barrier, either (III) just beneath the barrier, whereupon they continued to the interception point at the constant velocity described for the predictable targets, or (IV) at the interception point, where they flashed transiently. The time at which the unpredictable targets appeared beneath the barrier was made random by adding a jitter time (0-300ms) to the temporal delays of the predictable target conditions: 540+jitter time for condition III; 720+jitter time for condition IV. In unpredictable target conditions, the target onset times corresponded to the time during which the target was not visible. There was a distinction between the “predictable” and “unpredictable” conditions that participants could therefore infer from the initial context of each single trial. Specifically, if the target dropped toward the barrier, it always re-appeared just beneath the barrier at the specified time (consistent with target velocity), and then continued falling to reach the interception point, or suddenly flashed at the interception point at a specified time later. By contrast, if the target simply disappeared (without any motion) from the origin, then the subjects knew that the target onset time could not be predicted with certainty because of the inserted random delays.

**Figure 2:**
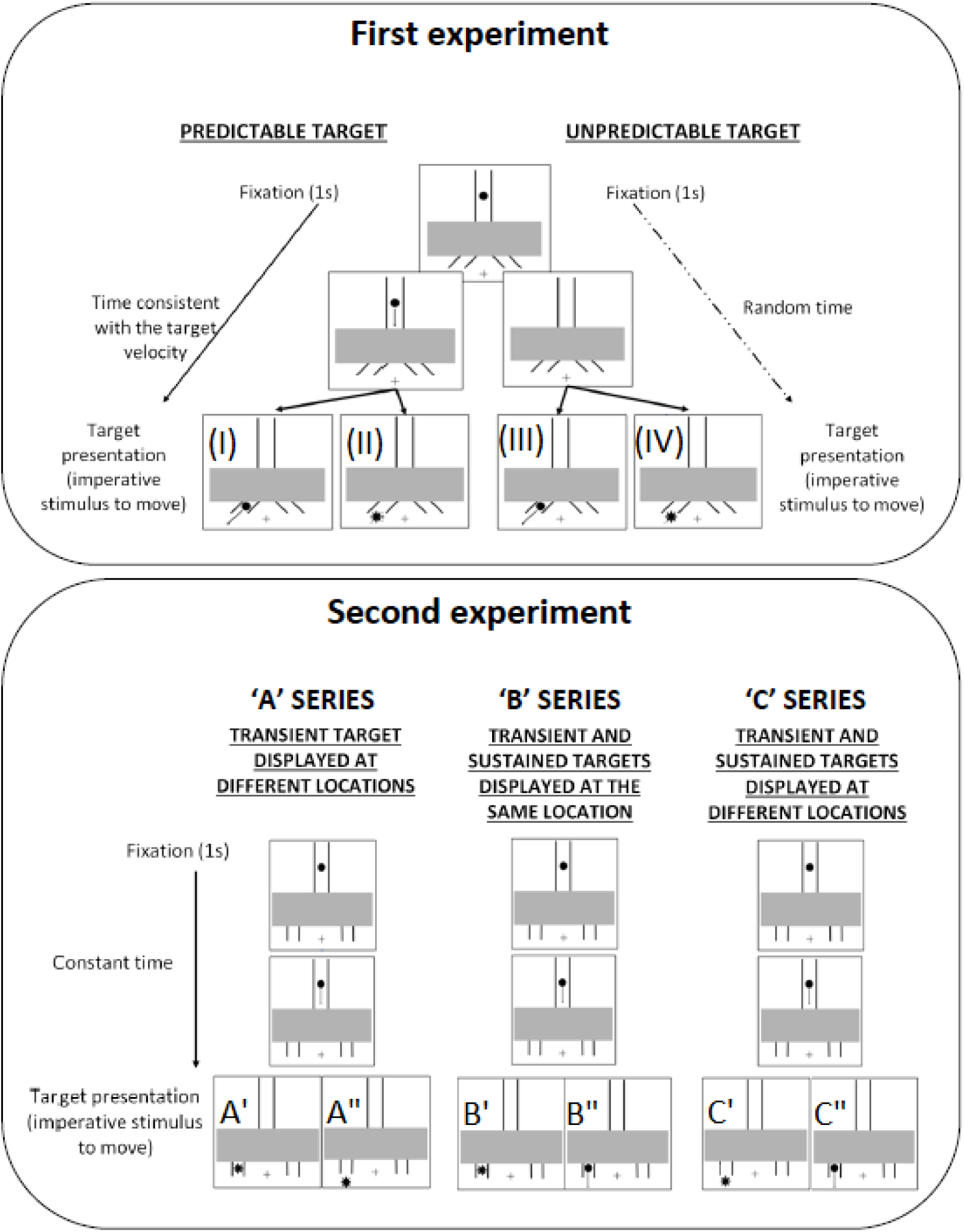
First experiment: timeline of the predictable and unpredictable targets. In the predictable target conditions, the stimuli dropped from the stem of the track at constant velocity of ∼35dva/s until it passed behind the barrier, and re-appeared either just beneath the barrier and continuing toward the interception point (I), or suddenly flashed at the interception point (II) at a time consistent with the target speed (i.e. ∼540ms for condition I; ∼720ms for condition II). In the unpredictable target conditions, the stimuli disappeared from the stem of the track and suddenly reappeared at a random time underneath the barrier, either just beneath the barrier (III), or at the interception point (IV), as described for the predictable targets. The onset time of the unpredictable targets was made random by adding a jitter time (0-300ms) to the temporal delays of the predictable target conditions (i.e. ∼540+jitter time for condition III; ∼720+jitter time for condition IV). Second experiment: timeline of the transient and sustained targets. In all target conditions, the stimuli dropped from the stem of the track at constant velocity of ∼35dva/s until they passed behind the barrier, and re-appeared underneath the barrier at ∼540ms from the start of the trial. In (A) series, the target appeared transiently just beneath the barrier (A’), or at the interception point (A”); in (B) series, the target appeared transiently just beneath the barrier (B’), or appeared just beneath the barrier and continued moving toward the interception point (B”); in (C) series, the target appeared transiently at the interception point (C’), or appeared just beneath the barrier and continued moving toward the interception point(C”). In both of the two experiments, the transient targets appeared by making one single flash of ∼8ms of duration.

The interception point was defined as the target trajectory point at which the hand could virtually reach the target, using only flexion/extension transverse plane movements of the shoulder. The participants could not bring the hand exactly to the physical target locations, hence veridical target interception was not achieved. The target always appeared as a full filled circle, thus avoiding a gradual emergence of the target from behind the barrier (rising moon stimulus), which would produce a high spatial frequency stimulus that has been reported to impair the SLR expression (Kozak et al. 2019). The participants were instructed to react as quickly as possible to the stimulus presentation, by moving the hand toward the virtual point of interception with the target. That is, they had to bring the hand to the interception point as soon as they saw the target, even when it appeared just beneath the barrier (i.e. conditions I and III, Figure 2).

To start the trial, participants were required to align the hand with the gaze fixation spot and to hold fixation for 1 second. They were instructed to stay relaxed as much as possible before the target appearance, and not to break fixation until the target re-appeared underneath the barrier to the left or right of the fixation point. On each trial, gaze-on-fixation was checked on-line with an EyeLink 1000 plus tower-mounted eye tracker device (SR Research Ltd., Ontario, Canada), at a sampling rate of 1000 Hz. If the fixation requirement was not respected, participants received an error message and the trial was repeated.

For this experiment, participants performed 10 blocks of 48 reaches each (480 total reaches). The trial-list design of each block was 2 (predictable or unpredictable target) x 2 (transient static or moving target upon reappearance) x 2 (left or right target location), so 8 unique trials types that were intermixed randomly in each block. As there were 10 blocks total, we obtained 60 repeats of every unique trial type.

#### Experiment 2: sustained vs transient stimulus

Experiment 2 was completed by 11 participants, nine of whom also participated in the first experiment. They performed visually guided reaches toward temporally predictable targets, following the same task rules described for experiment 1. In this experiment, the target was constrained to move within a track that was shaped as an inverted diapason, and it always started moving at a constant velocity before disappearing behind the barrier (Figure 2, bottom panel).

The participants performed reaches toward three different target types, which were compared pairwise in three separate series of 4 trial blocks with 60 reaches per block (12 total blocks with 60 reaches per block = 720 total reaches). The order of the three series was randomized across participants, and the task comparisons in each series were between:

- target appearing transiently just beneath the barrier, or target appearing transiently at the interception point (A series, figure 2 bottom panel);
- target appearing transiently just beneath the barrier, or target appearing just beneath the barrier and continuing toward the interception point (B series, figure 2 bottom panel);
- target appearing transiently at the interception point, or target appearing just beneath the barrier and continuing toward the interception point (C series, figure 2 bottom panel).

All targets dropped at constant velocity of ∼35dva/s until they passed behind the barrier, and constantly emerged from behind the barrier at ∼540ms from the start of the trial (onset time of target drop). In each target condition, the total disappearance time was ∼380ms. The eccentricity from the fixation spot was 10dva for both of the two target locations. The vertical distance between the ‘just beneath the barrier’ and ‘interception point’ spots was ∼6dva. The target always appeared below the barrier as a full, filled circle. In the transient target conditions the stimulus appeared with one flash of ∼8ms of duration.

### Data recording

Surface EMG (sEMG) activity was recorded from the clavicular head of the right pectoralis muscle (PMch), and from the posterior head of the right deltoid muscle (PD), with double-differential surface electrodes (Delsys Inc. Bagnoli-8 system, Boston, MA, USA). The quality of the signal was checked, using an oscilloscope, before starting the recording session. The sEMG signals were amplified by 1000, filtered with a 20-450Hz bandwidth filter, and full-wave rectified after digitization.

Arm motion was monitored by a three-axis accelerometer positioned flat on the lateral aspect of the right upper arm, in line with the right humerus, just proximal to the lateral humeral epicondyle. The sEMG and kinematic data were sampled at 2 kHz and stored on a computer using a 16-bit analog-digital converter (USB-6343-BNC DAQ device, National Instruments, Austin, TX, USA). Data synchronization was guaranteed by starting the recording of the entire data-set at the frame at which the target started moving (predictable conditions), or disappeared from the screen (unpredictable conditions).

The accelerometer enabled the determination of the point in time, relative to the stimulus presentation, at which the force produced by the muscle was enough to overcome the upper limb inertia and initiate direct arm movement; hereinafter this will be called reaction time (RT). Movement onset was discriminated with the use of the *cumulative sum* method (Basseville and Nikiforov 1993). More precisely, we defined the RT as the point in time at which the cumulative sum trace exceeded more than five standard deviations beyond the mean accelerometer signal, which was obtained by averaging the acceleration values recorded in the 100ms prior to target onset time. We excluded trials with RT<130ms (∼5% of the trials) as indicative of anticipation, as well as those with RTs>500ms as indicative of inattentiveness.

### Data analysis

#### Detection of SLRs

In line with previous approaches to quantify the SLR (Corneil et al. 2004; Pruszynski et al. 2010), we used a time-series ROC (receiver operator characteristic) analysis to identify the presence of an SLR. The ROC analysis indicates the probability that an ideal observer could discriminate the side of the stimulus location based solely on sEMG activity. Here, we used the ROC analysis to detect the point in time at which the location of the target can be discriminated (discrimination time, DT) from the sEMG trace. The DT provides the earliest available indication that the target location has been discriminated by some neural circuit in the brain, and that the motor command encoding this location has been delivered to the muscle. All analyses were aligned to the diode signal that detected the onset time of a secondary light displayed coincidently with the real target (see apparatus description).

For every muscle sample and tested condition, we separated the sEMG activity for all correct reaches based on visual stimulus location, and sorted the trials according to RT. Then we checked for an absence of voluntary muscle pre-activation by testing the correlation between the sEMG activity at target onset (during a 50ms window up to target presentation) and the RT. If a significant negative correlation was found, then we concluded that the muscle was pre-activated to react faster, thus potentially biasing the sEMG activity between the stimulus onset and the movement initiation, including the SLR epoch. In this case, no further analysis was conducted on the sEMG data (3.17% of the recordings were discarded due to muscle pre-target activation). For recordings not showing muscle pre-target activation, we compared the sEMG activity for the target requiring muscle activation (left target for the pectoralis; right target for the posterior deltoid) and the target requiring muscle inhibition (e.g. right target for the pectoralis; left target for the posterior deltoid). Muscle activity for the two targets was split into two equally-sized groups based on RT, subdividing fastest 50% of the trials (*fast* trial set) and the slowest 50% (*slow* trial set). We then conducted separate ROC analyses on both trial sets. We ran the ROC analysis on every data sample obtained between 100ms before and 300ms after the visual stimulus onset, and we calculated the area under the ROC curve (AUC). The AUC values range from 0 to 1, where a value of 0.5 indicates chance discrimination, whereas a value of 1 or 0 indicates perfectly correct or incorrect discrimination, respectively. We set the thresholds for discrimination at 0.65; this criterion exceeds the 95% confidence intervals of data randomly shuffled with a bootstrap procedure. The time of earliest discrimination was defined as the time after stimulus onset at which the AUC overcame the defined threshold, and remained above that threshold level for at least 15ms. In accordance with the literature, the candidate SLR was considered only if the discrimination time of fast and slow trial sets was between 80 and 120ms after visual stimulus onset (Gu et al. 2016).

In order to determine whether the short-latency muscle response was consistently time-locked to the visual stimulus onset, or co-varied with the RT, we compared the two DTs (fast trials DT; slow trials DT) for the average RT in the slow and fast trial sets by fitting a line to the data (Wood et al. 2015). If the slope of the line is 45°, then the discrimination time co-varies with RT, and so the initial sEMG is time-locked with the RT. Conversely, if the slope of the line is infinite (90°), then the DT remains the same irrespective of the RT, and the onset of sEMG is consistently linked to the appearance of the peripheral target. We classified an SLR observation as positive (+SLR) if the slope of the line was >67.5° (halfway between 45° and 90°). If the SLR was positively detected, then we ran a time-series ROC on *all* trials to determine the point in time at which the target location could be discriminated solely from the sEMG trace.

The DT variable that we extracted from the time-series ROC analysis is sensitive to the size of the muscle response to the stimulus, relative to the baseline sEMG trace. More precisely, the DT will be earlier for a strong muscle response than a weak one, even if both muscle responses deviate from background at the same time form the stimulus presentation. This is because the vigour of the muscular response to the visual stimuli directly influences how sharply the ROC curve rises toward the threshold defined for the discrimination time. In our experiment, this means that even if the ROC curves of two different target types (e.g. predictable, unpredictable) start diverging from chance (i.e. 0.5) at the same time after target onset, the location of the target eliciting the stronger responses can be discriminated earlier than the other one. This may mislead interpretation of the muscle response latency in the different task conditions. To provide a more sensitive determination of the visuomotor response onset time, we ran an analysis that searches for the inflection point at which the time-series ROC curve begins to deviate from chance toward the discrimination threshold (i.e. 0.65). To do this, we fit a DogLeg regression (Carroll et al. 2019; Pruszynski et al. 2008) to the ROC curve for each millisecond spanning from 50ms before target presentation up to the DT. The divergence time was then determined by taking the later time point between two possible candidates as the onset time of the response: (1) the point that minimizes the squared errors between the ROC curve and the DogLeg regression; (2) the last local minimum in the ROC curve before the DT. This analysis allowed us to reduce the influence of the visuomotor response size on the indexing of the time point at which the muscles started responding to the target.

#### Correlation of visual SLR magnitude with reaction time

In order to test the relationship between the +SLR and RT, we correlated RT with the magnitude of the EMG activity in the SLR time-window on a trial-by-trial basis (Pruszynski et al. 2010; Gu et al. 2016). More specifically, we defined the SLR magnitude as the mean sEMG activity recorded in the 10ms subsequent to the DT of the slow trial sets (see ROC method). We used this method to index the muscle activity in a time window restricted to a brief period consistent with the earliest time that the SLR-related EMG was present for both the fast and slow halves of the RT distributions. The aim was to minimise the potential for the index of SLR magnitude to be contaminated by EMG activity associated with the subsequent EMG burst that is time-locked to limb motion. Nonetheless, we also tested different methods to define the time window over which to quantify the SLR, including simple measurements of the peak and average activity within the nominal SLR time-window (80-120ms after stimulus presentation). Similar results were obtained irrespective of the specific method employed (i.e. negative correlation between the SLR magnitude and the RT; see results).

#### Statistical analysis

Statistical analyses were performed in SPSS (IBM SPSS Statistics for Windows, version 25, SPSS Inc., Chicago, Ill., USA). Results were analysed with one-sample, paired-sample and independent-sample T-Tests as the normality of the distributions was verified by the Shapiro-Wilk test. The Chi-squared test was used to analyse changes in SLR prevalence across predicable and unpredictable conditions. For all tests, the statistical significance was designated at *p*<0.05.

## RESULTS

In the raster plots that are shown in figure 3 (panels A and B), SLRs appear as a vertical band of either muscle activation (A) or inhibition (B) that is time locked ∼100ms to the stimulus onset time, and which only slightly co-varies with the voluntary RT. The consistency of the time-locking of the DT to the stimulus presentation was tested by running the ROC analysis over both fast and slow trial sets, and fitting a line connecting the two DTs with the average RT of the slow and fast sets (see materials and methods). When a +SLR was observed, the line slope exceeded 67.5°, meaning that the visuomotor response was more time-locked to the stimulus onset than to the RT (Figure 3D). By contrast, for the –SLR example the DT co-varied with the RT rather than being driven by the stimulus (Figure 3H).

**Figure 3:**
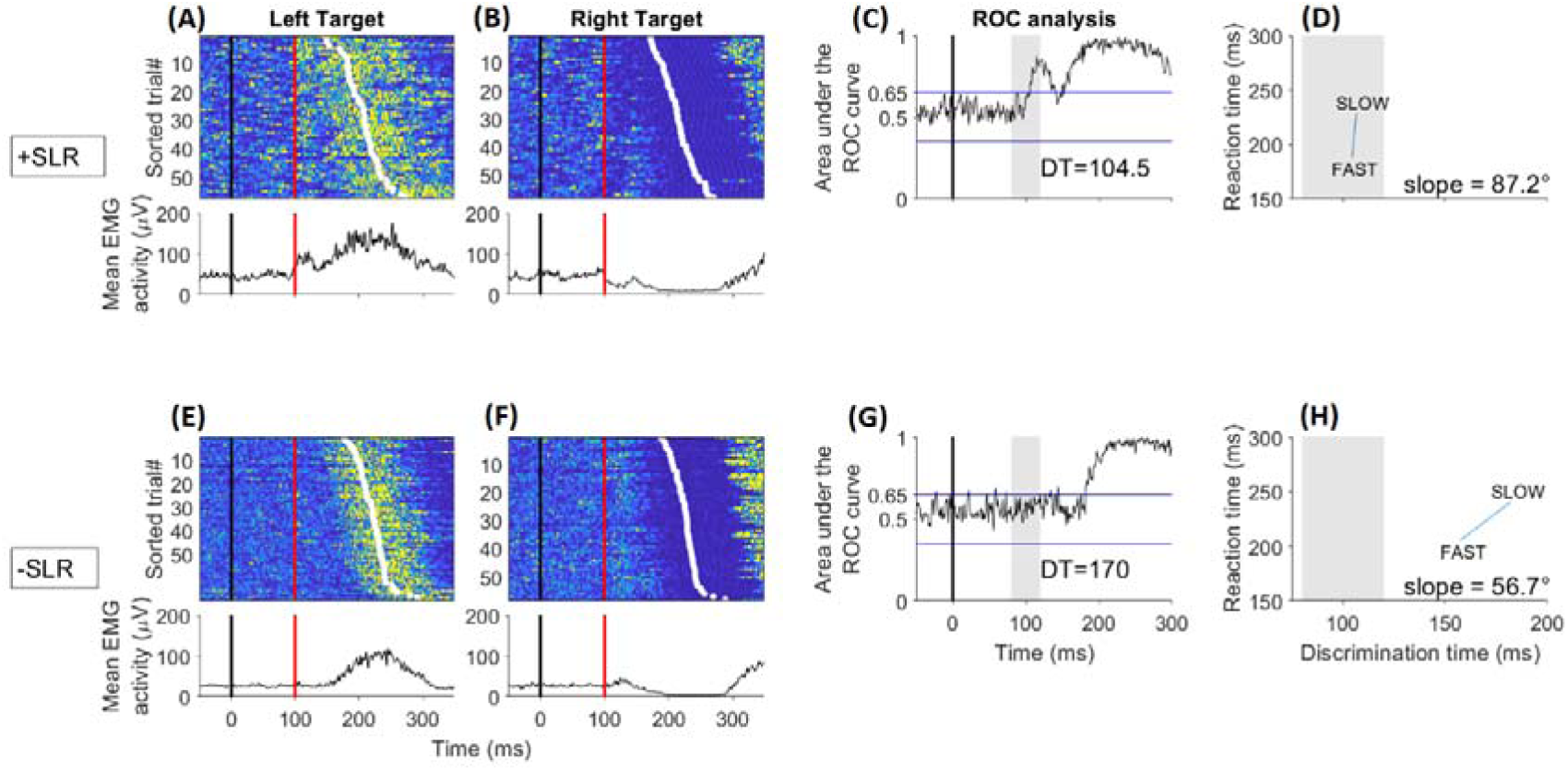
Surface EMG activity from the clavicular head of the pectoralis major muscle of an exemplar +SLR producer (subject 12, table 1) and an exemplar –SLR producer (subject 5, table 1) who participated in the first experiment. For both participants, the data are taken from to the trials in which the target appeared at a predictable time just beneath the barrier (condition I, figure 2). The muscle acts as agonist and antagonist for the left and right targets, respectively. For both subjects, rasters of rectified surface sEMG activity from individual trials are shown (darker yellow colours indicate greater sEMG activity; panels A, B, E, F), as are the traces of the mean sEMG activity. Data are aligned on visual target presentation (solid black vertical line at time 0) and sorted according to reaction time (white dots within the rasters). The solid red line indicates the expected initiation time of the SLR (∼100ms from target onset). The +SLR subject shows a column of either rapid muscle activation (A) or inhibition (B) time-locked to the stimulus onset regardless of the time of voluntary movement initiation. By contrast, for the –SLR subject, the muscle activity at 100ms from target presentation does not differ from the background level (panels E and F). The ROC analysis panels show the point in time (discrimination time -DT) at which the location of the target can be discriminated by the muscle activity. The DT is identified by the first time frame at which the area under the ROC curve surpasses the value of 0.65, and remains over this threshold for 15ms. For the +SLR subject, the discrimination time falls inside the SLR epoch highlighted by the grey patch (C), whereas the discrimination time of the –SLR producer exceeded the SLR time window (G). D and H panels show a line connecting the discrimination time identified by running the ROC analysis over the fast and slow trials. The two discrimination times are plotted for the slowest and fastest half of voluntary reaction times, and the line slope is showed. For the +SLR subject (D), both the early and late discrimination times are inside the SLR epoch evidenced by the grey patch, and the line slope exceeds 67.5°, thus indicating the presence of a visuomotor response that is more time-locked to the stimulus onset than to the reaction time. On the contrary, for the –SLR subject (H), the rapid visuomotor response is not observed and the line slope indicates that the onset of the movement-related sEMG response co-varies with the reaction time.

### Experiment 1 results

#### Predictable targets lead to more prevalent SLRs

In the first experiment, we investigated whether the effectiveness of the emerging target paradigm in eliciting SLRs relies on the predictability of the stimulus onset time. Trial-by-trial, the stimulus onset time was either predictable or unpredictable and, for each of the two predictability conditions, the target could appear either just beneath the barrier or at the interception point (see materials and methods).

**Table 1:** Occurrences of positive SLRs in the clavicular head of the pectoralis major muscle (PMch) and the posterior deltoid (PD) across participants in all four targets conditions tested in experiment 1.

| Subject | Predictable target conditions |  |  |  | Unpredictable target conditions |  |  |  |
| --- | --- | --- | --- | --- | --- | --- | --- | --- |
|  | Target appearing just beneath the barrier |  | Target appearing at the interception point |  | Target appearing just beneath the barrier |  | Target appearing at the interception point |  |
|  | PMch | PD | PMch | PD | PMch | PD | PMch | PD |
| 1 | x |  | x | + |  |  |  |  |
| 2 | x |  | x |  | x |  | x |  |
| 3 | x |  | x | + |  |  |  |  |
| 4 | x |  | x | + |  |  |  |  |
| 5 |  |  |  |  |  |  |  |  |
| 6 |  |  | x |  |  |  |  |  |
| 7 |  |  | x | + |  |  |  |  |
| 8 |  |  | x | + |  |  |  |  |
| 9 |  |  | x | + |  |  | x |  |
| 10 | x |  | x |  |  |  |  |  |
| 11 | x |  | x | + | x | + | x | + |
| 12 | x |  | x | + | x |  | x |  |
| 13 | x | + | x | + |  |  | x |  |
| 14 | x |  | x |  |  |  | x |  |
| 15 |  |  | x | + |  |  |  |  |
| Total +SLRs | 9 | 1 | 14 | 10 | 3 | 1 | 6 | 1 |

Table 1 reports whether or not SLRs were detected among all the subjects and conditions tested in the experiment 1. The table shows that SLRs were much more frequent for the trials in which the subject knew exactly when the target would appear compared to those in which the subject was cued that the timing would be random. For the PMch, SLRs were elicited in all but one subject when the predictably timed target appeared at the interception point, and in 9 of 15 subjects when the target appeared just beneath the barrier. For the PD, 10 subjects exhibited an SLR when the predictably timed target appeared at the interception point, but only one subject expressed an SLR when the predictably timed target appeared just beneath the barrier. SLRs were less prevalent but not entirely absent for the unpredictable timing condition. For the PMch, 6 subjects produced SLRs for targets at the interception point and 3 of them also exhibited an SLR when the predictable target appeared just beneath the barrier. For the PD, only one subject exhibited an SLR in both of the two unpredictably timed target conditions. Three subjects produced SLRs in the PMch under all four conditions. One subject produced no SLRs for any muscle or condition. The difference in SLR prevalence between the two muscle samples is likely due to the loading force, which led to increased baseline activity of the PMch while leaving unloaded the PD.

To determine the influence of temporal target predictability on the prevalence of SLR expression, we pooled the +SLR observations by selecting the participants who exhibited a +SLR with at least one of the two target locations beneath the barrier. For the PMch, 14 out of 15 participants expressed +SLRs with at least one of the two predictable target conditions, and 6 out of 15 participants exhibited +SLRs with at least one of the two unpredictable target conditions. These observations resulted in significantly higher prevalence of +SLR for predictable than unpredictable target conditions (chi-squared test; *p*=0.002, chi-squared= 9.6, df= 1; Figure 4). For the PD, we identified +SLRs in 10 out of 15 participants for the predictable target conditions, and 1 out of 15 participants for the unpredictable target conditions, again resulting in significantly higher prevalence of +SLR for predictable than unpredictable conditions (chi-squared test; *p*<10^−3^, chi-squared=12.9, df= 1; Figure 4).

**Figure 4:**
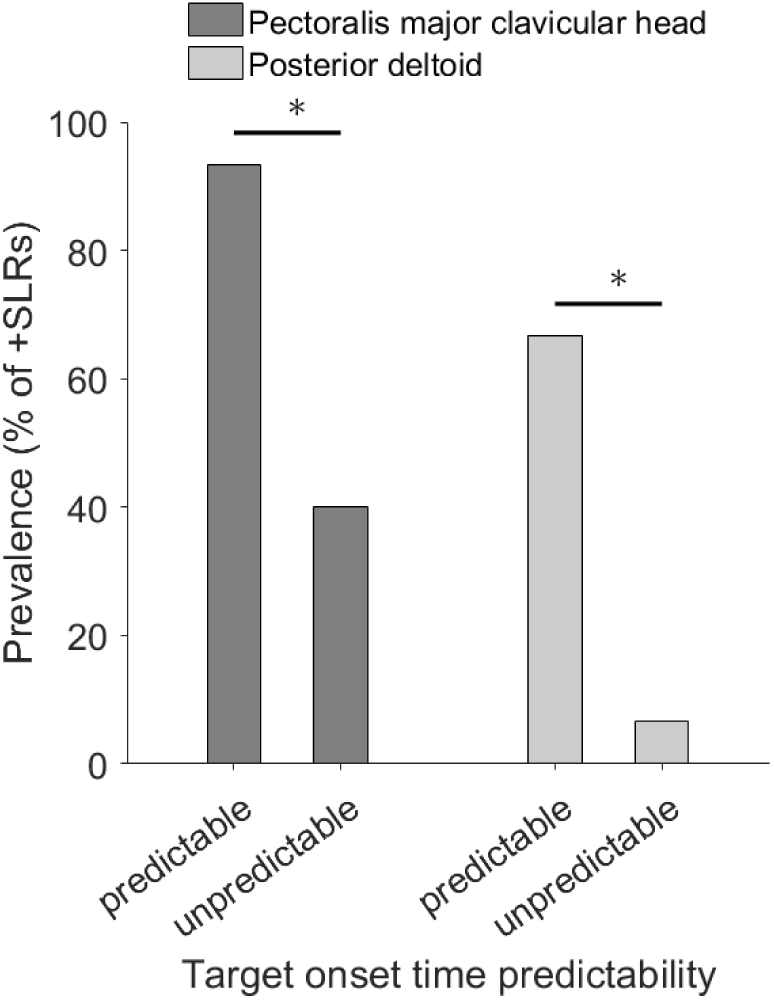
Dependence of +SLR prevalence on target onset time predictability. Significant differences between the predictable and unpredictable target conditions: * *p*< 0.01.

#### Predictable targets facilitate SLRs

To elucidate the effect of temporal predictability on the magnitude and timing of the SLR, we selected the participants who produced a +SLR under both predictably and unpredictably timed target conditions. For each participant meeting this criterion, if the SLR was detected with all the tested conditions (subjects 2, 11, 12; Table 1), we took the mean response onset time (DogLeg regression analysis), DT (ROC threshold analysis), and magnitude values (i.e. mean of the two predictable target conditions; mean of the two unpredictable target conditions). By contrast, if the number of +SLR observations mismatched between the temporal predictability conditions (subjects 13, 14; Table 1), we considered the values of the +SLR that was expressed to the same presentation spot for both predictable and unpredictable timed targets (i.e. interception point for subjects 13 and 14).

For the PMch, 6 out of 15 participants exhibited +SLRs with both predictable and unpredictable target conditions, whereas only one participant had +SLRs in both target predictability conditions for the PD. Given the low number of +SLR observations for the PD, we considered only data from the PMch sample to evaluate the effect of target onset predictability on the SLR.

Figure 5 shows the PMch activity for reaching movements toward predictable and unpredictable target conditions of an exemplar subject who participated in the first experiment, and who exhibited +SLRs to both predictable and unpredictable timed targets. The SLR magnitude was larger for the predictable than the unpredictable target condition (predictable target: 111.43μV; unpredictable target: 92.25μV). The ROC threshold analysis revealed that the target direction could be first reliably discriminated from the sEMG at 75.5ms for the predictable target (Figure 5C), and at 85.5ms for the unpredictable target (Figure 5C). However, the DogLeg regression analysis showed that the ROC curve began to deviate from chance at ∼70ms for both of the two predictability target conditions (Figure 5C).

**Figure 5:**
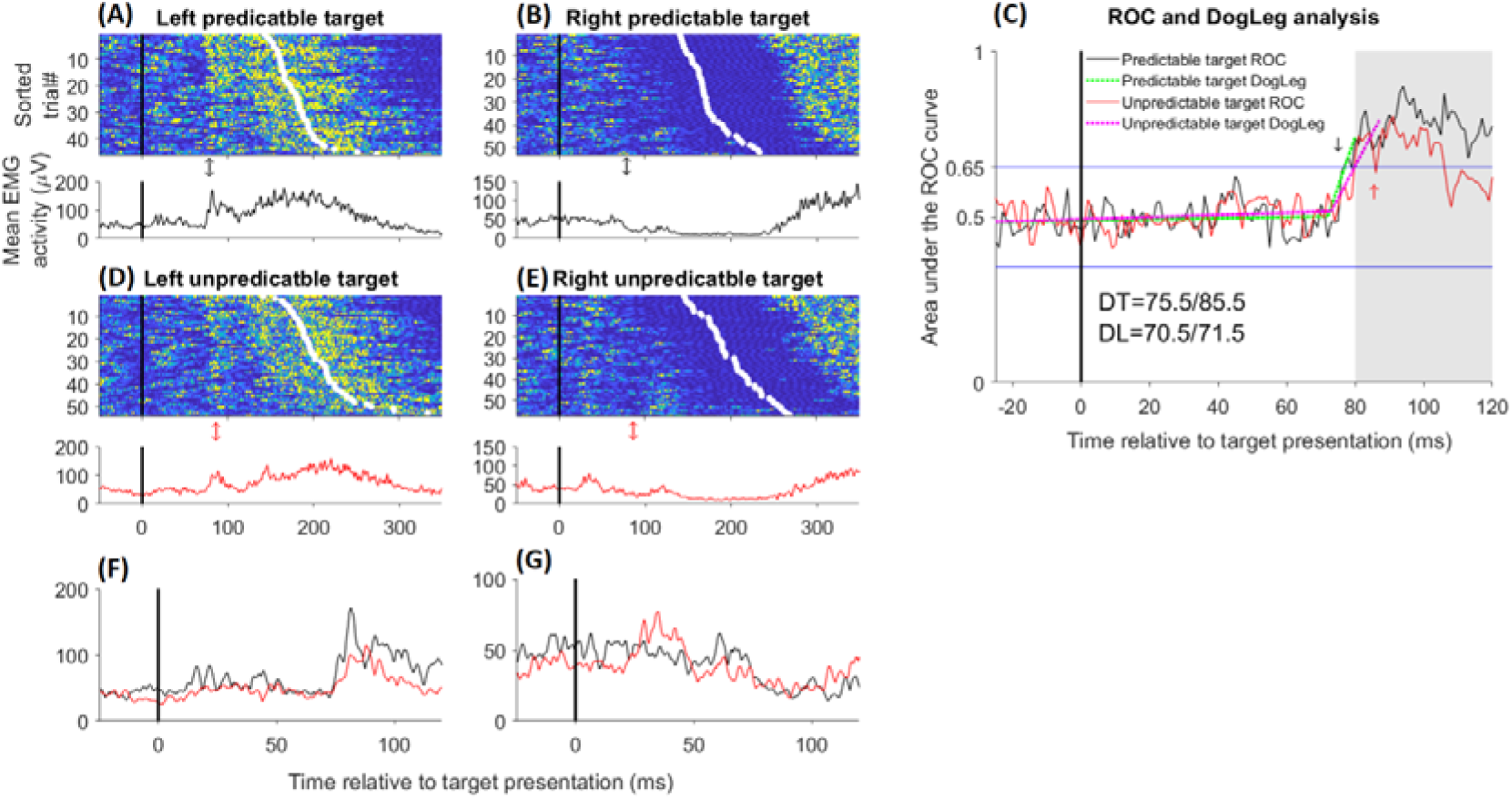
Surface EMG activity of the clavicular head of the pectoralis major muscle of an exemplar subject exhibiting +SLRs in both predictable and unpredictable target conditions. The data are from subject 12 (Table 1) in trials in which the target appeared transiently (1 flash of ∼8ms of duration) at the interception point. For both predictable and unpredictable targets, rasters of rectified surface sEMG activity from individual trials are shown (A, B, D, F; same format as figure 3), as is the trace of the mean sEMG activity. Panels F and G offer a zoomed view of the mean sEMG traces, and show how the SLR is larger with the predictable timed target (panel F, black trace) than the unpredictable timed target (panel F, red trace). However, the divergence onset time of the sEMG traces from baseline overlaps across the two temporal predictability conditions. C panel shows the results of the ROC analysis to identify the point in time (discrimination time - DT) at which the location of the target can be discriminated from the sEMG, and the results of the DogLeg (DL) regression to determine the onset time of the visuomotor response (see materials and methods). The DT of the predictable target condition is displayed as black arrows, whereas the DT of the unpredictable target condition is shown with red arrows. The ROC analysis reveals that the discrimination of the target location was earlier in predictable (75.5ms) than unpredictable (85.5ms) target conditions. However, the starting point of the ROC curve rising trend differs of just 1ms between the two temporal predictability conditions (predictably timed target, DL=70.5; unpredictably timed target, DL=71.5). That is, the difference in discrimination time between the predictably and unpredictably timed targets is not evident from the earliest initiation of the response to the stimulus, and is likely relative to the different magnitude of the short-latency response to temporal predictable and unpredictable targets.

The temporal predictability effects on the SLR timings and magnitude were consistent across the 6 participants exhibiting an SLR to at least one target location for both predictable and unpredictable timing. Specifically, the time at which the ROC curve started deviating from chance (DogLeg regression analysis) was not significantly different between temporal predictability conditions (Figure 6A), thus suggesting that the short-latency response to the target was initiated at the same time irrespective of target onset time predictability. By contrast, the discrimination time variable extrapolated from the ROC analysis showed that the location of the predictable target was discriminated significantly earlier in predictable than unpredictable target conditions (paired T-Test; t=-2.31, *p*=0.034; Figure 6B). Further, we observed significantly stronger SLRs for predictable than unpredictable targets (paired T-Test; t= 4.39, *p*=0.003; Figure 6C).

**Figure 6:**
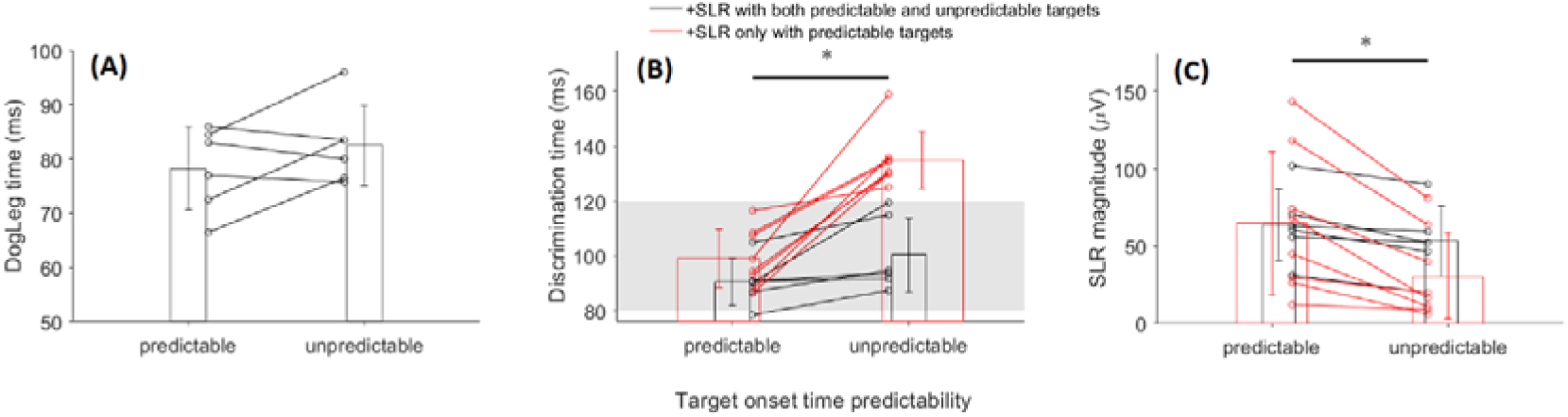
Latencies and magnitudes of the visuomotor responses on the PMch to stimulus presentation. Panel A shows when the area under the ROC curve started to deviate from chance, which indexed the onset time the visuomotor response to the target, via running a DogLeg regression analysis on the ROC trace (see materials and methods). Panel B shows when the area under the ROC curve exceeded the 0.65 threshold (see materials and methods) that identifies the point in time at which the location of the target can be discriminated from the muscle activity. Each black line represents one of the 6 subjects with +SLRs on both predictable and unpredictable targets, whereas each red line represents one of the 8 subjects exhibiting +SLRs only on predictable conditions. The ROC analysis (B) revealed that the location of predictable targets could be discriminated according to our threshold significantly earlier (* p< 0.05) than that of unpredictable targets. However, the divergence time at which the area under the ROC curve deviated from chance (A) was not significantly different across the temporal predictability conditions. Panel C shows the magnitude of the SLR encoding the location of the target that appeared to the left of the fixation spot, thus requiring the activation of the PMch. The predictable targets lead to significantly stronger (* p< 0.01) SLRs than unpredictable targets.

Eight subjects generated +SLRs only in predictable target conditions (see table 1). In those subjects, the target location could not be discriminated from the sEMG until ∼135ms after unpredictable stimulus presentation (solid red lines in figure 6B), well after the boundary defining the end of the SLR epoch (i.e. 120ms). Further, the magnitude of the SLR to predictable targets (defined as the mean sEMG activity recorded in the 10ms subsequent to the DT of the slow trial sets; see materials and methods) was much higher than the mean sEMG activity recorded from 80 to 120ms after unpredictable target presentation (Figure 6C, solid red lines; +SLR mean magnitude ∼70μV, - SLR mean magnitude ∼30μV; paired T-Test; t=4.43, *p*=0.001).

### Experiment 2 results: transient flashing targets facilitate fast and strong SLRs

In the second experiment, we investigated whether the location or temporal attributes (transient flash vs sustained motion) of the target influence the SLR when stimulus eccentricity and appearance time are matched. Nine of the 11 subjects who participated in this experiment also participated in the first experiment and expressed an SLR with at least one of the two predictably timed targets.

As is consistent with experiment 1, the emerging target paradigm facilitated the expression of SLRs. Indeed, all of the 11 subject exhibited an SLR on the PMch with at least one of the conditions tested in experiment 2 (Table 2). Again, SLRs were observed more frequently for the PMch than for the PD. For both muscle samples, the frequency of +SLRs was similar between the target conditions that were compared pairwise in each of the three series. Consistently, we did not observe any significant difference in +SLR prevalence between the target conditions tested in the second experiment (chi-squared test; PMch, A series *p=*1, B series *p*=0.138, C series *p*=0.062; PD, A series *p*=1, B series *p*=0.269, C series *p*=0.338).

**Table 2:**
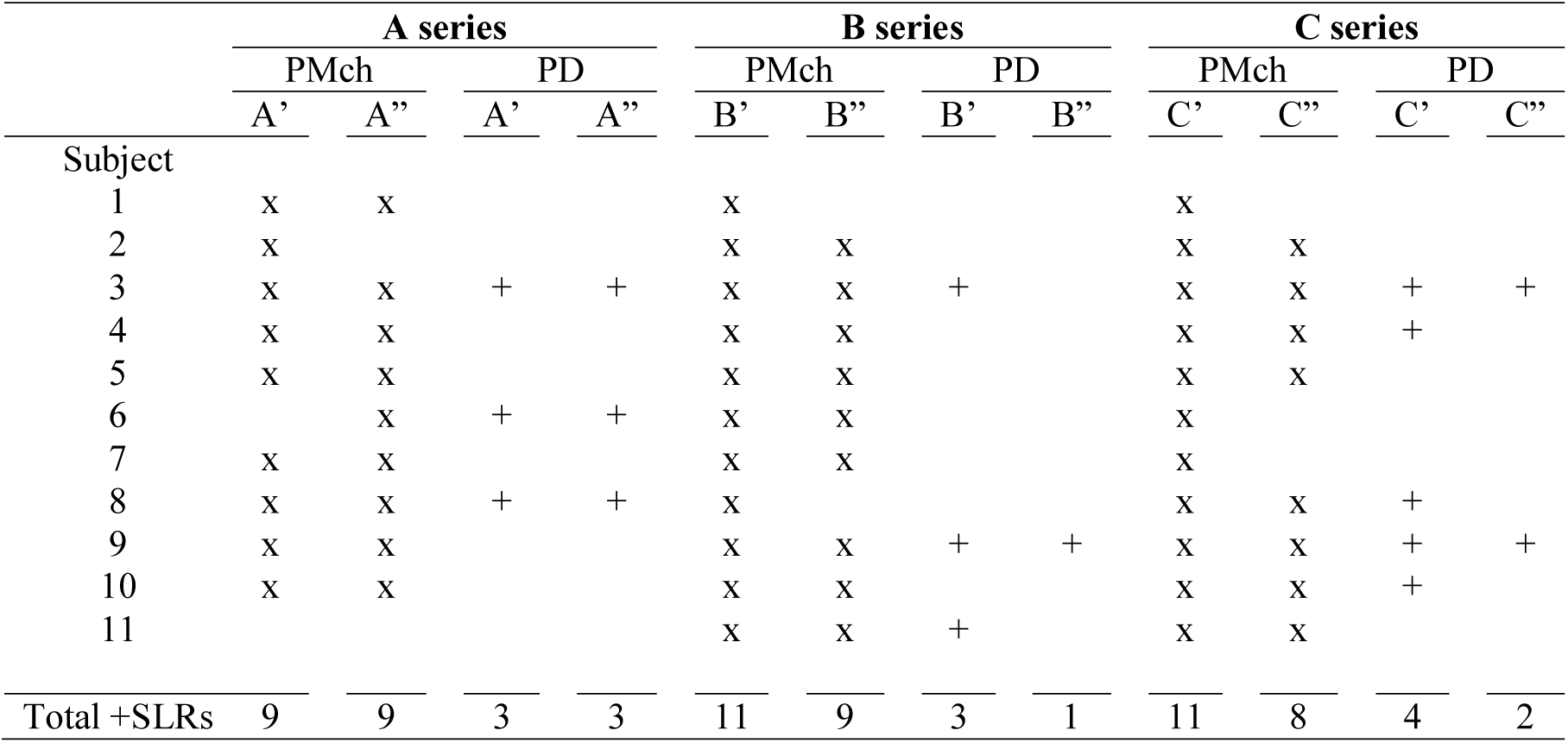
Occurrences of positive SLRs on the clavicular head of the pectoralis major muscle (PMch) and the posterior deltoid (PD) for each target condition tested in experiment 2. *A* series: comparison between the target that appeared transiently just beneath the barrier (A’), and the target that appeared transiently at the interception point (A”); *B* series: comparison between the target that appeared transiently just beneath the barrier (B’), and the sustained moving target that appeared just beneath the barrier and continued to the interception point (B”); *C* series: comparison between the target that appeared transiently at the interception point (C’), and the sustained moving target that appeared just beneath the barrier and continued to the interception point (C”). Subjects 1-9 correspond to subject 6, 2, 11, 9, 3, 10, 1, 13, and 12 in table 1.

|  | A series |  |  |  | B series |  |  |  | C series |  |  |  |
| --- | --- | --- | --- | --- | --- | --- | --- | --- | --- | --- | --- | --- |
|  | PMch |  | PD |  | PMch |  | PD |  | PMch |  | PD |  |
|  | A' | A'' | A' | A'' | B' | B'' | B' | B'' | C' | C'' | C' | C'' |
| Subject |  |  |  |  |  |  |  |  |  |  |  |  |
| 1 | x | x |  |  | x |  |  |  | x |  |  |  |
| 2 | x |  |  |  | x | x |  |  | x | x |  |  |
| 3 | x | x | + | + | x | x | + |  | x | x | + | + |
| 4 | x | x |  |  | x | x |  |  | x | x | + |  |
| 5 | x | x |  |  | x | x |  |  | x | x |  |  |
| 6 |  | x | + | + | x | x |  |  | x |  |  |  |
| 7 | x | x |  |  | x | x |  |  | x |  |  |  |
| 8 | x | x | + | + | x |  |  |  | x | x | + |  |
| 9 | x | x |  |  | x | x | + | + | x | x | + | + |
| 10 | x | x |  |  | x | x |  |  | x | x | + |  |
| 11 |  |  |  |  | x | x | + |  | x | x |  |  |
| Total +SLRs | 9 | 9 | 3 | 3 | 11 | 9 | 3 | 1 | 11 | 8 | 4 | 2 |

For each of the three series of pairwise comparisons (A, B and C series), we selected those participants who exhibited +SLRs for both of the two target conditions and compared the timings (DT, DogLeg) and magnitude of the SLRs. Again, a sufficient number of +SLRs to permit comparison was observed only for the PMch: *A* series, 8 and 3 subjects met this criterion for PMch and PD, respectively; *B* series, 9 and 1 subjects met this criterion for PMch and PD, respectively; *C* series, 8 and 2 subjects met this criterion for PMch and PD, respectively (Table 2).

Figure 7 shows the PMch activity of an exemplar subject who participated in the second experiment. The ROC analysis revealed similar DT for the transient flashing target presented at the two different locations beneath the barrier (A series). The time at which the ROC curve started to diverge from chance, as calculated by DogLeg regression, was also similar (A series, right panel). The magnitudes were also similar: 63.14μV for the target that appeared transiently just beneath the barrier; 58.82μV for the target that appeared transiently at the interception point. By contrast, SLR timing and magnitude differed between the sustained moving target and the transient target conditions, regardless of the location of the transient stimulus. More precisely, SLRs to the transient target could be discriminated above threshold at ∼20ms before the SLR to the sustained target (B and C series), and DogLeg regression showed that the ROC curve of the transient target started to deviate from chance more than 10ms before the ROC curve of the sustained target (right panels of B and C series). The sustained target SLR magnitude was also ∼25μV smaller than the SLR recorded with transient targets.

**Figure 7:**
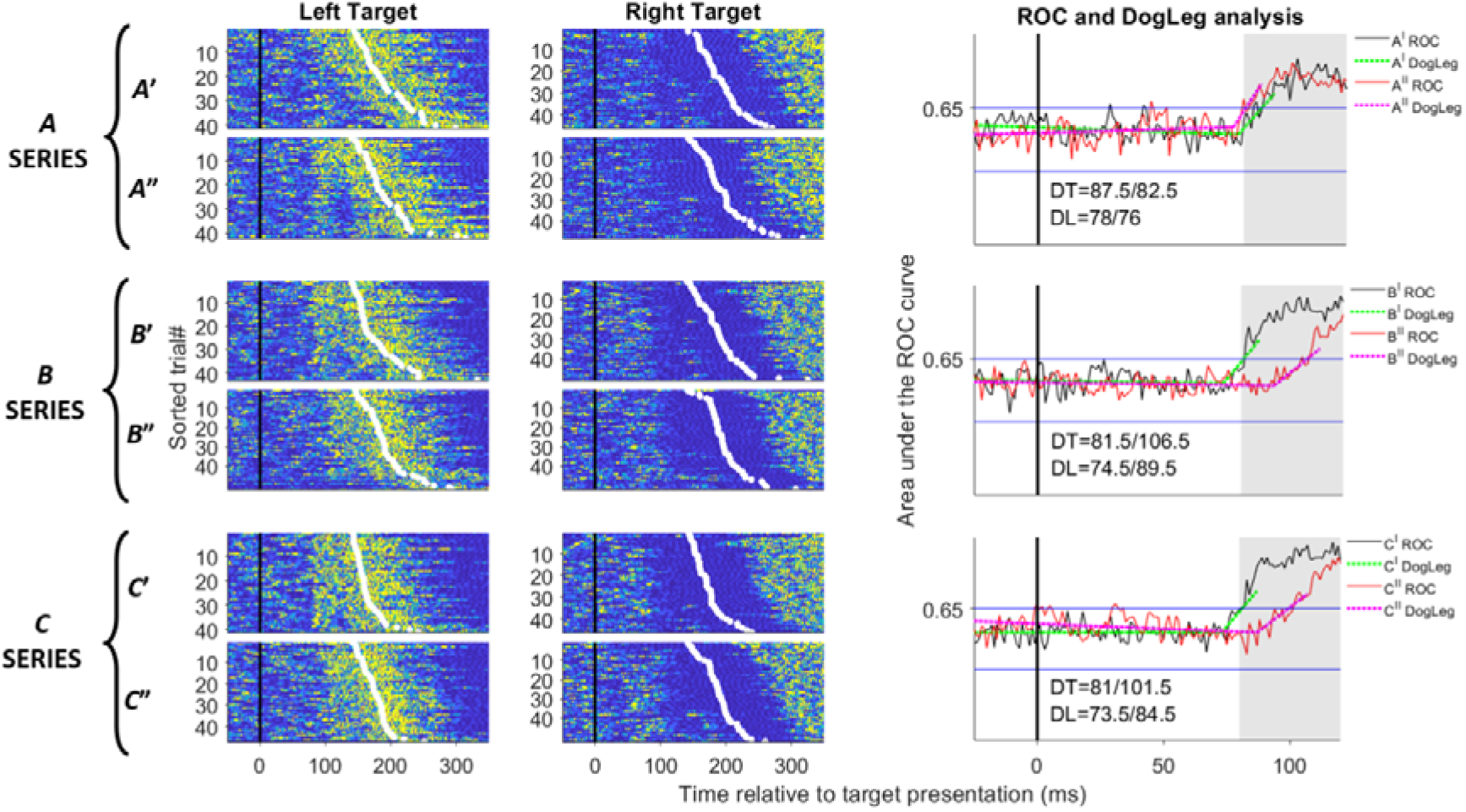
Surface EMG activity of the clavicular head of the pectoralis major muscle of an exemplar subject (9, Table 2) who participated in the second experiment. The panels show the three pairwise comparisons, as consistent with the design of the second experiment (see materials and methods, and Figure 2).For each target type, rasters of rectified surface sEMG activity from individual trials are shown (same format as figures 3 and 5). The ROC analysis reveals that the target location could be discriminated above threshold earlier for transient than sustained targets (right panels of B and C series). The different latencies of the visuomotor response to transient and sustained targets is also noticeable in that the column of the short-latency sEMG response is delayed in the rasters of the sustained target, with respect to the rasters of the transient targets. This accounts for the delayed initial deviation of the ROC curve from chance toward the discrimination threshold with the sustained targets, with respect to the transient targets (right panels of B and C series).

We observed similar trends across all participants of the second experiment. There were no statistically significant differences in the onset time of the visuomotor response (DogLeg time), discrimination time (DT), or SLR magnitude between the two transient targets that appeared at different distances from the bottom of the barrier (Figure 8, A series). Conversely, the SLR to transient stimuli started significantly earlier than to the sustained moving target (DogLeg time: paired T-Test, B series, t=3.36, *p*=0.005; C series, t=4.22, *p*=0.002; Figure 8), and also exceeded the discrimination threshold significantly earlier (discrimination time: paired T-Test, B series, t=16.99, *p*=<10^−3^; C series, t=7.8, *p*=<10^−3^; Figure 8). Further, the transient stimuli led to significantly stronger SLRs than sustained moving targets, regardless of transient stimulus location (paired T-Test, B series, t=-2.25, *p*=0.027; C series, t=-2.97, *p* =0.01; Figure 8).

**Figure 8:**
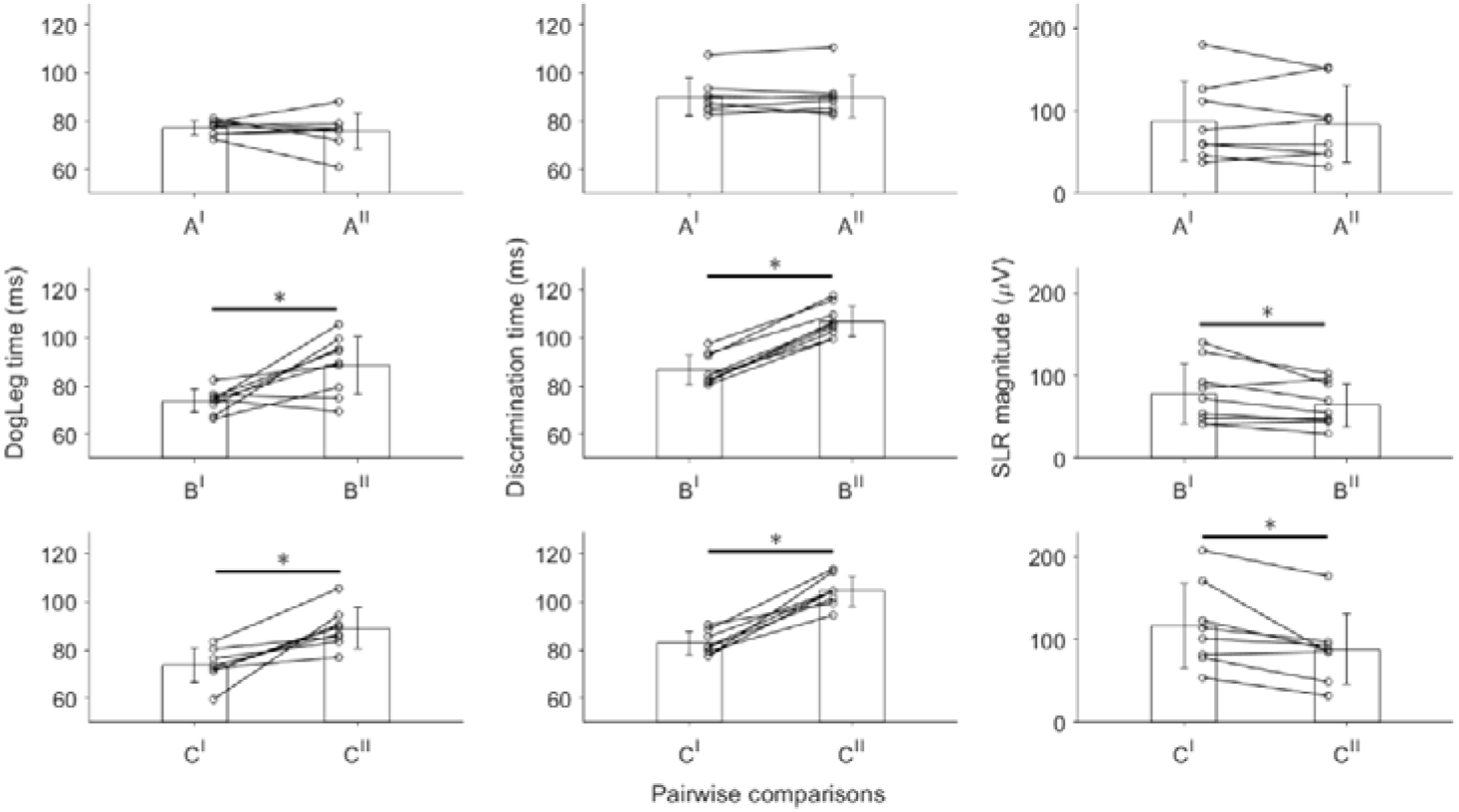
Latencies and magnitudes of the visuomotor responses to stimulus presentation. The DogLeg time represents the point in time at which the AUC starts to deviate from chance, whereas the discrimination time represents the point in time at which the target location could be discriminated from the sEMG activity above the threshold in the ROC area under the curve (see materials and methods). Each solid black line represents one subject having +SLRs on both target types used for the pairwise contrasts, as consistent with the design of the second experiment (see materials and methods, and Figure 2). The first row of panels shows the absence of significant differences between the transient target appearing just beneath the barrier (A’) and the transient target appearing at the interception point (A”). The second and third rows show that the transient target (B’ and C’) led to significantly faster (** *p*<10-3) and stronger (* *p*< 0.05) SLRs than the sustained targets (B” and C”), regardless of the location of the transient targets (B’ transient target just beneath the barrier; C’ transient target at the interception point).

### SLR magnitude correlates with the latency of the voluntary movement initiation

In both experiments, the SLR magnitude correlated with RT. The relationship for a single exemplar subject and correlation coefficients across all subjects are illustrated in Figure 9. In the first experiment, the SLR was negatively correlated with RT for both the predictable and unpredictable targets (one sample T-Test; predictable targets, t = −9.95, *p*<10^− 3^; unpredictable targets, t = −7.4, *p*<10^−3^). In the second experiment, the SLR magnitude was negatively correlated with RT for all three target presentation conditions (one sample T-Test; transient target just beneath the barrier, t = −6.86, *p*<10^−3^; transient target at the interception point, t = −6.81, *p*<10^−3^; sustained target, t = −5.52, *p*<10^−3^). That is, the latency of the RT tends to become longer as the magnitude of the SLR decreases, and vice versa.

**Figure 9:**
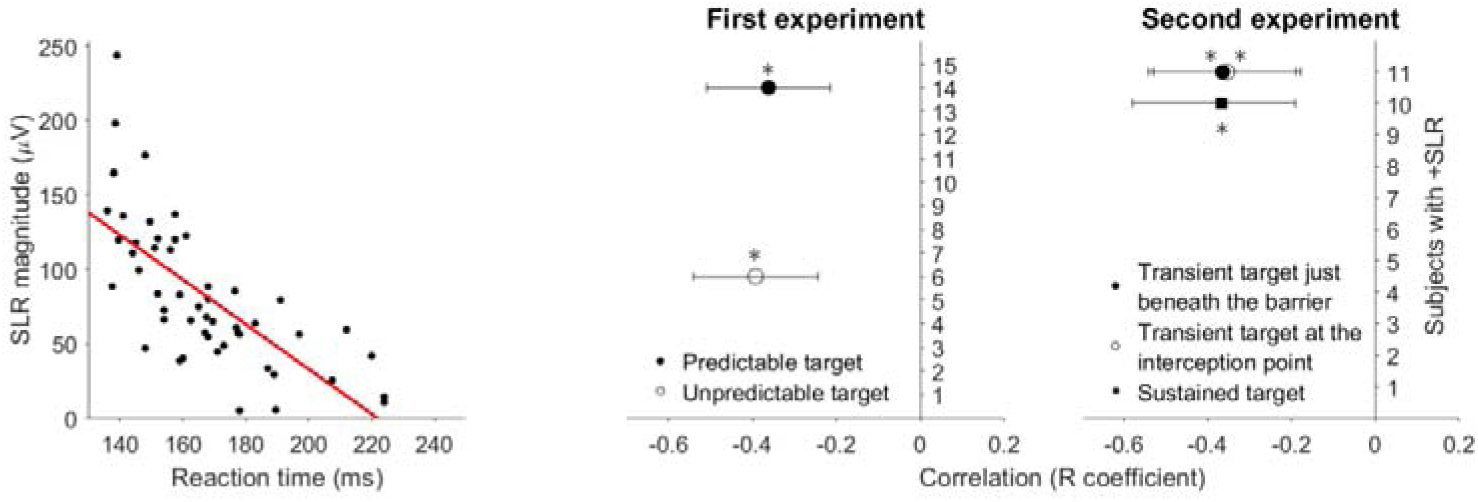
First panel shows the correlation between the reaction time and SLR magnitude from the pectoralis major clavicular head for an exemplar participant. Each data point represents a single trial and the solid red line is the linear regression function. The second and third panels show the average correlation coefficient of all participants who exhibited an SLR in experiment two. Regardless of target onset predictability (first experiment) and target temporal attributes (second experiment), the magnitude of the SLR demonstrates a significant negative correlation (* p<10-3) with the movement initiation.

## DISCUSSION

### Methodological factors in SLR prevalence

The distinctive short latency of the SLR suggests that it is a behavioural marker for the contribution of a subcortical system to the expression of rapid visuomotor behaviours. However, because early reports found such rapid responses to be sporadic across subject and conditions, the SLR has mainly remained a curiosity. Methods to consistently record robust SLRs across participants are therefore needed to facilitate research on this phenomenon, including whether or not it can be exploited for practical applications (e.g. training, rehabilitation). One objective of the current study was to identify experimental conditions that can generate robust SLRs in most subjects. Notably, Kozak et al. (2020) detected SLRs in all five of the participants that they tested, by adopting a moving target paradigm involving a visual barrier to partially occlude the target trajectory of motion. Here, we observed +SLRs among all but one of the 21 unique subjects tested with different versions of the moving target paradigm. This suggests that the low SLR prevalence that was previously reported (see Pruszynski et al. 2010) is not due to an absence of the neural circuitry necessary to generate an SLR, but rather to the use of stimulus paradigms that were suboptimal for the relevant neural pathway. Indeed, a negative SLR producer in one paradigm may become a +SLR producer in another (e.g. the 8 subjects that exhibited SLRs only with predicably timed targets, table 1, figure 6). We expect that most humans can be primed to produce SLRs given appropriate task conditions. This would imply that a subcortical system for visuomotor transformations is a ubiquitous feature of human sensorimotor control systems, which should be considered as a potential contributor in general theories of human motor behaviour.

Our experimental setup had some key differences from that adopted by Kozak et al. (2020). In that study, the authors presented the paradigm via a horizontal mirror that reflected a down-facing monitor, which also projected a real-time cursor to provide the participants with the visual feedback of the hand location. This resulted in a veridical spatial representation of both the hand and the target and allowed the participants to bring the hand exactly to the physical target locations. By contrast, we projected our stimulus on a monitor located in front of the participants and we limited their arm movements to the transverse plane, irrespective of the target location in the vertical axis. Indeed, the participants did not move their hand toward the exact target positions. Despite these differences, we replicated the high (>90%) SLR detection rate that was reported by Kozak et al. (2020). Thus, the emerging target paradigm may be a powerful tool to elicit consistent and robust SLRs, irrespective of minor differences in setup. Like Kozak et al. (2020), we recorded the muscle activity via surface EMG electrodes. The emerging target paradigm might therefore help to broaden the investigation of SLRs among populations that may be less tolerant of intramuscular electrodes (e.g. young, old and clinical populations).

It is worth noting that our experimental paradigm explored only conditions with tonic activation of the shoulder flexor muscles, including the PMch, against an extensor load. This may account for the large asymmetry of agonist-antagonist SLR production reported here and suggests further experiments to study the effects of direction and magnitude of such preloads.

### Neural mechanisms of SLR generation

In the first experiment, we investigated whether the SLR-facilitation effect of the emerging target paradigm relies on the temporal predictability of the stimulus presentation. We observed that the expression of SLRs in the emerging target paradigm is facilitated when the target onset time is predictable. The positive effects of stimulus onset time predictability on the expression of rapid visuomotor behaviours is consistent with earlier work that used the gap task paradigm (i.e. constant time gap between the warning stimulus and the imperative stimulus; Fischer and Boch 1983; Pruszynski et al. 2010; Wood et al. 2015; Glover and Baker 2019). Pruszynski et al. (2010) observed only a ∼44% of +SLR prevalence with the gap task, whereas the emerging moving target paradigm led to 100% SLR detection score in the work by Kozak et al. (2020), and above 90% in the present investigation, at least for the PMch sample. This suggests that the internal timing mechanism necessary to anticipate a temporally predictable stimulus may be facilitated if the target timing information is conveyed through a moving stimulus, rather than by the offset of a static fixation spot as in the gap task paradigm (Pruszynski et al. 2010; Wood et al. 2015; Glover and Baker 2019). This conclusion remains tentative, however, because we did not test predictable target conditions in which the target timing information is provided by extinguishing a static target in this study. Future work should compare the ‘original’ emerging target paradigm with a task akin to our unpredictable target conditions, but with a constant time between the offset of the target and its re-emergence underneath the barrier. This would provide a direct test to disentangle the effective influence of moving and static warning stimuli on the SLR expression when the target onset time is matched.

The SLR implies the existence of a short-latency neural pathway that quickly connects the retinal information with limb muscles, thus producing rapid visuomotor transformations. This pathway has been proposed to include the superior colliculus and its projections to the reticular formation, which in turn is connected with spinal interneurons and motoneurons (Pruszynski et al. 2010; Wood et al. 2015; Gu et al. 2016; Gu et al. 2018; Gu et al. 2019; Atsma et al. 2018; Kozak et al. 2019; Glover and Baker 2019). We propose that the baseline activity in superior colliculus neurons can be enhanced by prediction and motor preparation signals, likely originating from frontal and parietal cortical areas that project to superior colliculus (Boehnke and Munoz 2008; Dash et al., 2018). Such known cortico-collicular projections might raise the activity of the collicular neurons closer to threshold level, thus facilitating a visually evoked SLR from the retino-tectal or retino-geniculo-cortico-tectal pathways. This idea is supported by previous work showing that the generation of an express saccade is predicted by the pre-stimulus firing rates of superior colliculus neurons, which can be enhanced by the predictability of visual stimulus onset time (Dorris et al. 1997, 2002). Decrements in the production of express saccades in non-human primates were observed consistently in association with reduced collicular pre-stimulus preparatory activity, which was induced by transiently inactivating the frontal eye field area (Dash et al, 2018).

Considering that both express saccades and SLRs are thought to rely on the activity of neurons in the intermediate layers of the superior colliculus, we propose that target onset time predictability facilitates the expression of SLRs through a top-down priming of the superior colliculus. The assumption of top-down modulation of the SLR-system is consistent with previous work from Gu et al. (2016), who showed that the magnitude of the SLR was lower in the anti-reach than pro-reach tasks. This suggests that the task-context influences the state of the SLR-circuitry and the relative vigour of the rapid visuomotor response in muscles, potentially via cortical top-down modulation of the superior colliculus. Here, we add to this literature by showing that the expression of an SLR is facilitated when the stimulus onset time can be predicted from contextual information that must be interpreted within each trial, consistent with continuous cortical modulation of the superior colliculus.

In the second experiment, we compared the SLR to a transient flashing target that appeared just beneath the barrier with a transient flashing target that was presented at the interception point (see materials and methods; A series, figure 2). We found that the SLR did not depend on the location of the transient target in the vertical axis, at least within the range of vertical visual angles (∼6dva) explored in this experiment. Broader ranges of vertical distance between the target emerging spots should be investigated in future studies in order to understand the spatial resolution of a priming pathway. Trial-by-trial, the target moved at a constant velocity (∼35dva/s) before disappearing behind the barrier, and emerged (transiently) at a constant time (∼540ms) from the trial start. If the implied motion of the target behind the barrier was extrapolated assuming a constant target velocity, then it should have enabled the temporal prediction of the target that appeared just beneath the barrier. By contrast, the timing predictability of the target that appeared at the interception point would require the extrapolation of a target that seems to start accelerating while it is hidden by the barrier. In our setup, the appearance location of the target was randomized trial-by-trial, thus making impossible for the participants to know in advance the final location of the target and, thereby the implied target kinematics (acceleration = 0 or acceleration ≠ 0) from which to extrapolate its implied motion behind the barrier. The absence of significant target location-induced effects on SLR expression suggests that implied target motion per-se is not necessary to facilitate the SLR. More likely, the disappearance of the moving target behind the barrier enables more precise initiation of the internal timing signal than simply extinguishing the target, which would be consistent with the results of the first experiment. A cortical origin for this hypothetical internal timing signal seems likely but has not been specifically addressed here and needs further elucidation.

The previously reported effects of a moving target for facilitating SLRs might have been the result of its kinematic-related salience as a stimulus instead of (or in addition to) its utility for extrapolating timing information. Another question is whether the SLR circuitry responds differently to a target that suddenly appears and starts moving, and to a static target that flashes briefly. In both experiments, we compared responses to sustained moving targets and transiently flashed targets that appeared at predictable times. Previous work suggests that the visual neurons of the mammalian superior colliculus are sensitive to the kinematic features of visual stimuli. An fMRI study showed that the presentation of a cluster of dots, which were equally distributed around a fixation spot, enhanced the response of the human superficial collicular layer when the dots changed their state from static to dynamic (radial movements toward, or away from, the fixation spot; Schneider and Kastner 2005). Another fMRI study showed that the activity of the rat superior colliculus neurons was larger when a moving stimulus passed through their receptive visual field than when a static stimulus appeared within their receptive field (Lau et al. 2011). Noteworthy, earlier work also showed that the superficial layer of the human superior colliculus responds well to transient stimuli (Schneider and Kastner 2005). More precisely, Schneider and Kastner (2005) reported larger collicular responses with a flickering stimulus than with a moving stimulus. Further, Chen and Hafed (2018) observed that the non-human primate superior colliculus neurons have preference for low flicker stimulus frequencies (e.g. 3-10Hz). Specifically, the collicular neurons responded to low flicker frequencies with transient increments in firing rate (>160 spikes/s) encoding each individual flicker cycle. By contrast, higher stimulus flickering frequencies resulted in a less transient and more sustained response characterized by low-frequency firing rates (∼40 spikes/s), especially when the stimulus approached the flicker fusion frequency for perception (i.e. ∼60Hz of flickering frequency; Chen and Hafed 2018). This suggests that the superior colliculus is sensitive to the duration of the visual stimulus. The temporal events of visual stimuli are encoded by different cell types located in the superficial layers of the mammalian superior colliculus: ‘On’, ‘Off’, ‘On-Off’ cells (Wang et al. 2010). Notably, the On-Off cells outnumber the other cell types and their receptive fields overlap those of the other cells (Wang et al. 2010), suggesting a preferential superior colliculus representation for transient visual stimuli.

In the first experiment, we successfully detected SLRs in 14 people when the transient target appeared via a single flash of 8ms at the interception point, and in nine people when the sustained moving target appeared just underneath the barrier (PMch, table 1). This is consistent with a stronger effect from transience than motion, but contextual features of the paradigm might also have contributed to this result. Specifically, the subjects knew that if the target did not appear just beneath the barrier at a time consistent with the target velocity (540ms from the initial target drop), then it could only appear transiently at the lower location. The SLR facilitation observed with the transient target conditions could therefore be due to spatial predictability of the target re-emerging location. In the second experiment, we made the target location unpredictable from the trial-context by matching the timing of the transient and sustained moving targets. The more powerful and earlier SLRs with the transient target suggests that its effectiveness is related to it transient nature rather than the spatial predictability of the target location.

The target that appeared transiently should have triggered a synchronized and high-frequency response (Chen and Hafed 2018) to both the onset and the offset (8ms later) phases of the target, and in all the cells that can encode these temporal events (i.e. On, Off and On-Off cells; Wang et al. 2010). The strong and rapid transient collicular responses should result in the generation of high-frequency trains of action potentials whose spatiotemporal integration would facilitate rapid transmission through the projecting neurons of the brainstem reticular formation and spinal interneurons required to reach the motoneurons. By contrast, the detection of a moving target might require a longer time to encode its kinematics features by spatial integration across adjacent portions of retina and superior colliculus receptive fields. This could account for the delayed (∼10-20ms) SLR expression observed with the sustained moving targets as compared to the static transient targets (see second experiment results). Further, given its extended “on” phase, the sustained moving target would be encoded by the subpopulation of superior collicular *On* cells (Wang et al. 2010), including a low-frequency response (Chen and Hafed 2018). This should result in a weaker SLR as compared with a transient stimulus that could rapidly synchronize high-frequency responses (Chen and Hafed 2018) in multiple subpopulations of collicular cells (On, Off and On-Off cells; Wang et al. 2010), as consistent with the results from the second experiment. From an ethological standpoint, transient stimuli might trigger particularly brisk responses to orient the eyes, head, and limbs toward targets that may soon become unavailable. This correlates with the primitive function of the superior colliculus to acquire unknown but salient visual stimuli (Boehnke and Munoz 2008). Nonetheless, we are mindful that further studies are needed to elucidate the effective timing and intensity of the collicular visual response in these behavioural paradigms.

### Behavioural function of SLRs

A key outstanding question about the SLR concerns its functional role in motor behaviour. The term “stimulus-locked response” was originally coined by Pruszynski et al. (2010) to describe short latency muscle responses that were more invariant to the time of stimulus presentation than to movement initiation. We acknowledge that the timing of the SLR is variable trial-by-trial; similar variability is also seen in the first spike timing of visual responses in the intermediate superior colliculus (Peel et al. 2017). Regardless, we caution that the term SLR, or similar terms such as “rapid visuomotor responses” (Glover and Baker 2019) or “visual responses on muscles” (Corneil et al. 2004) may incorrectly impute a sensory rather than motor function. The selective muscle activation or inhibition that is tuned correctly to the direction of the reach would be expected to contribute the reach mechanics. We replicated previous correlation analyses (Pruszynski et al. 2010; Gu et al. 2016) to show that the magnitude of the SLR predicts the onset time of the voluntary movement. For every subject exhibiting a +SLR, we observed a significant negative correlation between SLR magnitude and RT indicating that stronger SLRs were associated with quicker mechanical responses.

Mechanistically, large and rapid activation of an agonist muscle and inhibition of a tonically active antagonist will shorten the time at which sufficient net torque is produced to overcome limb inertia and start accelerating the arm (i.e. RT). During the so-called long-latency voluntary phase of the reaching movement, the time needed by the rising force to start accelerating the arm will be reduced if the same motor units have already fired once during the earlier SLR. When the interval between action potentials in a muscle fiber is substantially shorter than the typical inter-spike interval for voluntary muscle recruitment (a phenomenon known as doublets; Van Cutsem et al. 1998), the total muscle contractile activation so-produced is greatly accelerated and sustained as a consequence of calcium diffusion and reuptake kinetics in the muscle fibres, a phenomenon called the *catch property* of muscle (originally reported by Burke et al. 1970 and reviewed mechanistically in Tsianos and Loeb 2017). If the same motor units participate in both the SLR and the voluntary reaction, the enhanced force output would be expected to contribute to a shorter RT. This could be resolved with intramuscular recording of discriminable, single-unit EMG signatures such as employed by Van Cutsem et al. 1998 to identify doublets in human muscles.

### Conclusions

Our results suggest that the effectiveness of the emerging target paradigm for eliciting SLRs is related to the temporal predictability of target presentation. Predictability of the stimulus onset time appears to prime the putative subcortical circuit responsible for SLRs, potentially via the generation of an internal timing signal. A plausible source of the timing signal is the descending cortico-cullicular projections, suggesting a top-down modulation of the SLR-circuitry. The fact that transient targets are especially effective in prompting SLRs is consistent with the known sensitivity of the superior colliculus to the onset and offset phases of transient visual stimuli. The effectiveness of the emerging target paradigm for facilitating consistent and robust SLRs greatly enhances the capacity of researchers to investigate the neural processes underling these express visuomotor responses, as well as their potential use in clinical and sporting applications.

## Acknowledgements

This work was supported by operating grants from the Australian Research Council (DP170101500) awarded to T.J. Carroll, B.D. Corneil, G.E. Loeb and G. Wallis.

